# Activation of Multiple Eph Receptors on Neuronal Membranes Correlates with The Onset of Traumatic Optic Neuropathy

**DOI:** 10.1101/2023.06.05.543735

**Authors:** Thomas A. Strong, Juan Esquivel, Qikai Wang, Paul J. Ledon, Hua Wang, Gabriel Gaidosh, David Tse, Daniel Pelaez

## Abstract

**Background:** Optic neuropathy (ON) is a major cause of irreversible blindness, yet the molecular determinants that contribute to neuronal demise have not been fully elucidated. Several studies have identified ‘ephrin signaling’ as one of the most dysregulated pathways in the early pathophysiology of ON with varied etiologies. Developmentally, gradients in ephrin signaling coordinate retinotopic mapping via repulsive modulation of cytoskeletal dynamics in neuronal membranes. Little is known about the role ephrin signaling played in the post-natal visual system and its correlation with the onset of optic neuropathy.

**Methods:** Postnatal mouse retinas were collected for mass spectrometry analysis for Eph receptors. Optic nerve crush (ONC) model was employed to induce optic neuropathy, and proteomic changes during the acute phase of neuropathic onset were analyzed. Confocal and super-resolution microscopy determined the cellular localization of activated Eph receptors after ONC injury. Eph receptor inhibitors assessed the neuroprotective effect of ephrin signaling modulation.

**Results:** Mass spectrometry revealed expression of seven Eph receptors (EphA2, A4, A5, B1, B2, B3, and B6) in postnatal mouse retinal tissue. Immunoblotting analysis indicated a significant increase in phosphorylation of these Eph receptors 48 hours after ONC. Confocal microscopy demonstrated the presence of both subclasses of Eph receptors in the inner retinal layers. STORM super-resolution imaging combined with optimal transport colocalization analysis revealed a significant co-localization of activated Eph receptors with injured neuronal processes, compared to uninjured neuronal and/or injured glial cells, 48 hours post-ONC. Eph receptor inhibitors displayed notable neuroprotective effects after 6 days of ONC injury.

**Conclusions:** Our findings demonstrate the functional presence of diverse Eph receptors in the postnatal mammalian retina, capable of modulating multiple biological processes. Pan-Eph receptor activation contributes to the onset of neuropathy in ONs, with preferential activation of Eph receptors on neuronal processes in the inner retina following optic nerve injury. Notably, Eph receptor activation precedes neuronal loss. We observed neuroprotective effects upon inhibiting Eph receptors. Our study highlights the importance of investigating this repulsive pathway in early optic neuropathies and provides a comprehensive characterization of the receptors present in the developed retina of mice, relevant to both homeostasis and disease processes.

## INTRODUCTION

Neuropathic diseases of the retina are a leading cause of irreversible blindness worldwide [1]. While many risk factors associated with the development of neuropathic diseases are known, the molecular determinants for the onset of synaptic instability, neurite retraction, and subsequent neuronal loss are yet to be fully elucidated. Identification of appropriate molecular targets is a crucial step in the development of effective therapies that can halt or reverse neuropathic progression and preserve useful sight. Pan-genomic and pan-proteomic profiling of glaucomatous and traumatic optic neuropathies has allowed for the construction of the first-ever roadmaps for system-wide characterization at the molecular level for the pathophysiology of common neuropathic mechanisms [2–5]. Strikingly, most of these studies have revealed that ‘ephrin signaling’ is one of the most dysregulated signaling cascades in the neurodegenerative process. Studies have further shown that signaling via activated Eph receptors in early neuropathic states is evident in both animal models and human samples. Eph receptors (Eph) and ephrin ligands (efn) constitute the largest family of receptor tyrosine kinases in mammalian biology. To date, 16 Eph receptors have been identified and divided into subfamily A and subfamily B based on their sequence homologies and binding affinities [6–10].

Developmentally, Eph/efn signaling plays an important role in axon guidance and topographic mapping of neuronal projections [11–16], synaptogenesis and dendritic spine morphology [17–24], and synaptic plasticity and remodeling [19,25–27]. Eph/efn signaling is distinctive in that its signal is transduced bidirectionally, with receptor-ligand interactions initiating signaling cascades in both receptor-expressing (forward) as well as ligand-expressing (reverse) cells simultaneously. Reverse signaling (efn-mediated) is generally regarded as an attractive and stabilizing stimulus for neuronal extensions and stabilizing synaptic connections [28,29], whereas activated forward signaling (Eph-mediated) is repulsive to outgrowing neurites [7,30] and is responsible for inducing axonal growth cone collapse [31–37], restricting mid-line crossing of axons [38–42], and destabilizing synaptic connections [42–46]. During development, Eph/efn signaling is responsible for establishing guiding gradients that direct the retinotopic projections of RGC axons onto the visual centers of the brain [11–14,47–51]. This guidance for the outgrowing RGC neurites established by Eph/efn signaling is achieved through graded repulsion rather than attraction [7,31,32,38]. The upregulation of developmentally relevant programs in adult tissue is generally associated with mechanisms for repair and healing. However, the anachronic activation of growth repulsive pathways such as Eph forward signaling could result in detrimental outcomes. In fact, Eph receptor engagement and activation has been associated with several neurodegenerative disease states of the central nervous system (CNS) including Alzheimer’s disease [44,45,52–55]; glaucomatous degeneration of the retina [2–5,56–61]; traumatic brain injury [62–64]; stroke [65–67]; and spinal cord injury [68–71].

The CNS lacks the ability to undergo endogenous repair and given the repulsive nature of Eph receptor signaling, it is imperative to determine the role of these repulsive guidance programs in the neurodegenerative process. To this end, the identification of the molecular targets involved is the first step towards developing effective therapies for these diseases. In this study, we identified the Eph receptors expressed in the postnatal mouse retina, evaluated their early temporal expression and activation after injury, and determined their localization within the retinal layers and on specific cellular compartments following injury. We advance the hypothesis that neuropathic onset involves, at least in part, the reactivation of repulsive ephrin forward signaling on retinal neuronal membranes based on our findings. These results provide a rationale to evaluate modulation of this signaling pathway as a novel treatment for the management of optic neuropathies.

## MATERIALS AND METHODS

### Optic Nerve Crush Model (ONC)

All experiments involving mice were carried out in accordance with the ARVO statement for the Use of Animals in Ophthalmic and Vision research and were approved by the Animal Care and Use Committee at the University of Miami. Optic nerve crush (ONC) injury was performed on either male C57BL/6J WT or male Thy1-GFP mice at 2 months of age. The ONC procedure was performed as previously described [72]. Briefly, all mice were anesthetized via intraperitoneal injection of ketamine (80 mg/kg) / xylazine (10 mg/kg); their optic nerves were exposed intraorbitally and crushed with jeweler’s forceps (Dumon #5; tip dimension, 0.1 x 0.6 mm) for 10 seconds approximately 1 to 2 mm behind the optic disk, pupillary response using indirect illumination from the side was used to indicate a successful injury. Animals were allowed to recover from anesthesia and maintained in standard housing conditions for the duration of the specified experimental time points. Animals were euthanized and perfused with 1x phosphate buffered saline (PBS) (Corning, Cat.# 21-040-CV), after which the eyes were enucleated, and retinas micro-dissected. Dissected retinas were either fixed in 4% formaldehyde (MilliporeSigma, Cat.# FX0415-4) in PBS for 2 hours at room temperature or lysed with RIPA buffer (Thermo Fisher Scientific, Cat.# 89900) containing protease and phosphatase inhibitors (Thermo Fisher Scientific, Cat.# A32961) depending on downstream analysis requirements.

### Mass Spectrometry

Proteomic analysis was done on dissected retinal tissue of male C57BL/6J WT mice at 14 days (*N* = 5 biological replicates, pooled), 2 months (*N* = 5 biological replicates, pooled), and 12 months of age (*N* = 4 biological replicates, pooled). Samples were processed and analyzed by solution digestion and 90-minute data-independent acquisition by MS Bioworks Protein Mass Spectrometry Service, Ann Arbor, MI, USA. Briefly, flash frozen dissected retinal tissue was lysed in a modified RIPA buffer (50 mM Trish HCl, pH 8.0, 150 mM NaCl, 2.0% SDS, 0.1% TX100, 1x Roche Complete Protease Inhibitor) using 1.4 mm stainless steel beads in a Next Advance Bullet Blender, 2 cycles x 3 minutes each. Samples were then heated to 60°C for 30 minute and centrifuged at 16,000 x g. Each sample was then TCA precipitated overnight at -20°C, pellets were washed and resuspended in 8 M urea, 50 mM Tris HCL, pH 8.0, 1x Roche Complete Protease Inhibitor. An equal aliquot of each pooled sample (14 days, 2 months, and 12 months) was taken to create a combined sample. 50 mg of combined sample was digested overnight with trypsin. Samples were reduced for 1 hour at room temperature in 12 mM DTT followed by alkylation for 1 hour at room temperature in 15 mM iodoacetamide. Trypsin was added to an enzyme:substrate ratio of 1:20. Each sample was acidified to 0.3% TFA and subjected to SPE using Waters mHLB.

DIA chromatogram library generation was done using 1 mg of the pool and analyzed by nano LC/MS with a Water M-class HPLC system interfaced to a ThermoFisher Exploris 480. Peptides were loaded on a trapping column and eluted over a 75 mm analytical column at 350 nL/min; both columns were packed with XSelect CSH C18 resin; the trapping column contained a 5 mm particle; the analytical column contained a 2.4 mm particle. The column was heated to 55°C using a column heater. A 90-minute gradient was employed. The mass spectrometer was operated in data-independent mode. Six gas-phase fractions injections were acquired for 6 ranges: 396 to 502, 496 to 602, 596 to 702, 696 to 802, 796 to 902, and 896 to 1002. Sequentially, full scan MS data (60.000 FWHM resolution) was followed by 26 x 4 m/z precursor isolation windows, another full scan and 26 x 4 m/z windows staggered by 2 m/z; products were acquired at 30,000 FWHM resolution. The automatic gain control target was set to 1e6 for both full MS and product ion data. The maximum ion inject time was set to 50 milliseconds for full MS and “dynamic” mode for products with 9 data points required across the peak; the NCE was set to 30. 1 mg per sample was injected at random and analyzed by nano LC/MS with a Waters M-class HPLC system interfaced to a ThermoFisher Exploris 480. Peptides were loaded on a trapping column and eluted over a 75 mm analytical column at 350 nL/min; both columns were packed with XSelect CSH C18 resin; the trapping column contained a 5 mm particle; the analytical column contained a 2.4 mm particle. The column was heated to 55°C using a column heater. A 90-minute gradient was employed. The mass spectrometer was operated in data-independent mode. Sequentially, full scan MS data (60,000 FWHM resolution) from m/z 385-1015 was followed by 61 x 10 m/z precursor isolation windows, another full scan from m/z 385-1015 was followed by 61 c 10 m/z windows staggered by 5 m/z; products were acquired at 15,00 FWHM resolution. The maximum ion inject time was set to 50 milliseconds for full MS and “dynamic” mode for products with 9 data points required across the peak; the NCE was set to 30. An injection of the sample pool was included at the start, middle, and end of the batch. DIA data were analyzed using Scaffold DIA 3.2.1.

### Western Blots

Retinal tissues (*N* = 3 biological replicates) were lysed with RIPA buffer (Thermo Fisher Scientific, Cat.# 89900) containing protease and phosphatase inhibitors (Thermo Fisher Scientific, Cat.# A32961), and protein concentrations were measured using the DC Assay (Bio-Rad Laboratories, Cat.# 5000114), according to the manufacturer’s protocol. An equal amount of protein samples was loaded and separated on an SDS-PAGE 7.5% PROTEAN TGX Stain-Free gel (Bio-Rad, Cat.# 4568024), then transferred to 0.2 µm PVDF membranes (Bio-Rad Laboratories, Cat.# 1704156). The PVDF membranes were blocked in 5% Nonfat dry milk (Bio-Rad Laboratories, Cat.# 1706404) in Tris-buffered saline with 0.1% Tween 20 (TBST) (VWR, Cat.# K873-4L). Samples were probed with the primary antibodies listed in Supplemental Table 1 overnight in 5% bovine serum albumin (Gold Biotechnology, Cat.# A-420-100) in TBST. Blots were incubated with horseradish peroxidase-conjugated species-specific secondary antibodies listed in Supplemental Table 1. Proteins were visualized with an enhanced chemiluminescence substrate (Thermo Fisher Scientific, Cat.# 34095) and digitally imaged on a ChemiDoc MP Imaging System (Bio-Rad Laboratories)

### Immunofluorescent Staining for Confocal Microscopy and Stochastic Optical Reconstruction Microscopy (STORM)

Dissected ONC and uninjured WT retinal tissue from C57BL/6J was fixed for 2 hours at room temperature in 4% formaldehyde (MilliporeSigma, Cat. # FX0415-4) in PBS. ONC and uninjured tissues were embedded in 4% low-melting agarose (IBI Scientific, Cat.# IB70051) in PBS and sectioned (50 µm) using a Leica VT1000 S Vibrating blade microtome (Leica Biosystems). The sectioned tissues were permeabilized for 20 minutes in PBS containing 0.3% Triton X-100 (Thermo Scientific, Cat.# 85111) and blocked for 1 hour in PBS containing 10% normal donkey serum (Abcam, Cat.# ab7475). Primary antibodies (listed in Supplemental Table 3) in PBS solution were applied overnight at 4°C. AlexaFlour secondary antibodies (listed in Supplemental Table 2) were applied for 2 hours at room temperature. We used 4’, 6-diamidino- 2-phenylindole (DAPI) (1 ug/mL; Bio-Rad, Cat.# 1351303) to counterstain for confocal microscopy. The sectioned tissues were mounted on microscope slides with ProLong Diamond antifade mountant (Thermo Fisher Scientific, Cat.# P36965) Dissected ONC and uninjured retinal tissue from Thy1-GFP was fixed for 2 hours at room temperature in 4% formaldehyde (MilliporeSigma, Cat.# FX0415-4) in PBS. Flat-mount preparations were mounted on microscope slides with ProLong Glass antifade mountant (Thermo Fisher Scientific, Cat.# P36982).

### Confocal Microscopy Imaging

Confocal imaging was performed on a Leica AOBS SP8 confocal microscope (Leica Microsystems, Exton, PA). A HC PL APO 40x/1.30 OIL CS2 objective lens was used and imaged with a continuously adjustable galvo scanner. Fluorescence-labelled proteins were excited by 405, 488, and 561 nm lasers. Three conventional PMTs and one high sensitivity PMTs (HyD) were utilized to capture optical signals. Image acquisition and processing were accomplished on LAS X software.

### Stochastic Optical Reconstruction Microscopy (STORM) Imaging

STORM imaging and processing TIRF Imaging experiments were done with a Nikon eclipse Ti2 inverted microscope equipped with Nikon Instruments (N-STORM). A 100x TIRF objective 1.49NA objective lens was utilized and imaged using a Hamamatsu C11440 ORCA-flash CMOS 4.0 camera. Images were acquired sequentially 10,000 frames per filter channel at 20 millisecond time duration. Retinal tissue (*N* = 3 biological replicates) was labeled with JF646 secondary (Supplemental Table 2) were excited with 90% laser power from a 647 nm laser and A568 secondary (Supplemental Table 2) labeled samples were excited with a 561 nm laser at 100% laser power. Nikon Nd2 files were separated and converted to tiff files per channel by custom python script. STORM localization analysis was carried out with either ImageJ, thunderstorm plugin (1.3-2014-11-08) or WindSTORM MATLAB code. Data was fitted with Gaussian PSF model using weighted least-squares estimation for thunderstorm plugin.

### Optimal Transport Colocalization Analysis (OTC)

Individual regions of interest were processed using the OTC package [73] in R 4.2.1 by taking five 64 x 64 random samples with a matching counterpart for the second channel (Supplemental Figure 1). The OTC curves were then paired and compared using the Mann-Whitney U test.

### Intravitreal injection of Eph Receptor Inhibitors

Thy1-GFP mice were first anesthetized via intraperitoneal injection of ketamine (80 mg/kg)/xylazine (10 mg/kg). A quantity of 2 μL of Eph receptor inhibitor (Supplemental Table 3) was then injected into the temporal part of one eye via a glass micropipette inserted just behind the ora serrata (intravitraitreal injection). All small molecules were dissolved in Dimethy sulfoxide (DMSO) (Thermo Fisher Scientific, Cat.# D12345). All animals were treated immediately after ONC injury and again 48 hours post ONC injury.

### Sholl Analysis

Individual RGC images taken from the confocal microscope were imported and analyzed in FIJI, ImageJ2 (Version: 2.9.0/1.53t) software with the neuroanatomy, Sholl Analysis plug-in. The program creates concentric circles from 10 to 200 µm, each with a radius of 5 μm, starting from the soma and outwards towards the dendritic branching. Total length and branching were analyzed for each image [74,75].

### Statistical Analysis

One-way ANOVA and Mann-Whitney U tests were calculated using GraphPad Prism 9 (San Diego, CA), with a *p* value of *p* < 0.05 considered statistical significance.

## RESULTS

### Eph Receptor Analysis in Postnatal Retinas

Until now, it was unclear which of the many Eph receptors expressed during retinotopic development were still present in postnatal retinas. We used Mass Spectrometry Data Independent Acquisition (MS-DIA) to identify Eph receptors present in the uninjured postnatal retina in 14-day-old, 2-month-old, and 1-year-old C57BL/6J mice. Our data showed that of the 16 known Eph receptors, 3 class-A receptors (EphA2, EphA4, and EphA5) and 4 class-B receptors (EphB1, EphB2, EphB3, and EphB6) are present in the postnatal mouse retina (Figure 1), and that their relative quantities remain unchanged throughout the lifespan of the animals. Of the expressed receptors, EphB2 and EphA4 were measured to be the most abundant, whereas EphA2 is the least common in all age groups (Figure 1). Based on our MS-DIA data we narrowed our focus from the 16 known Eph receptors [76] to the detected 7 receptors in subsequent experiments.

**Figure 1.**
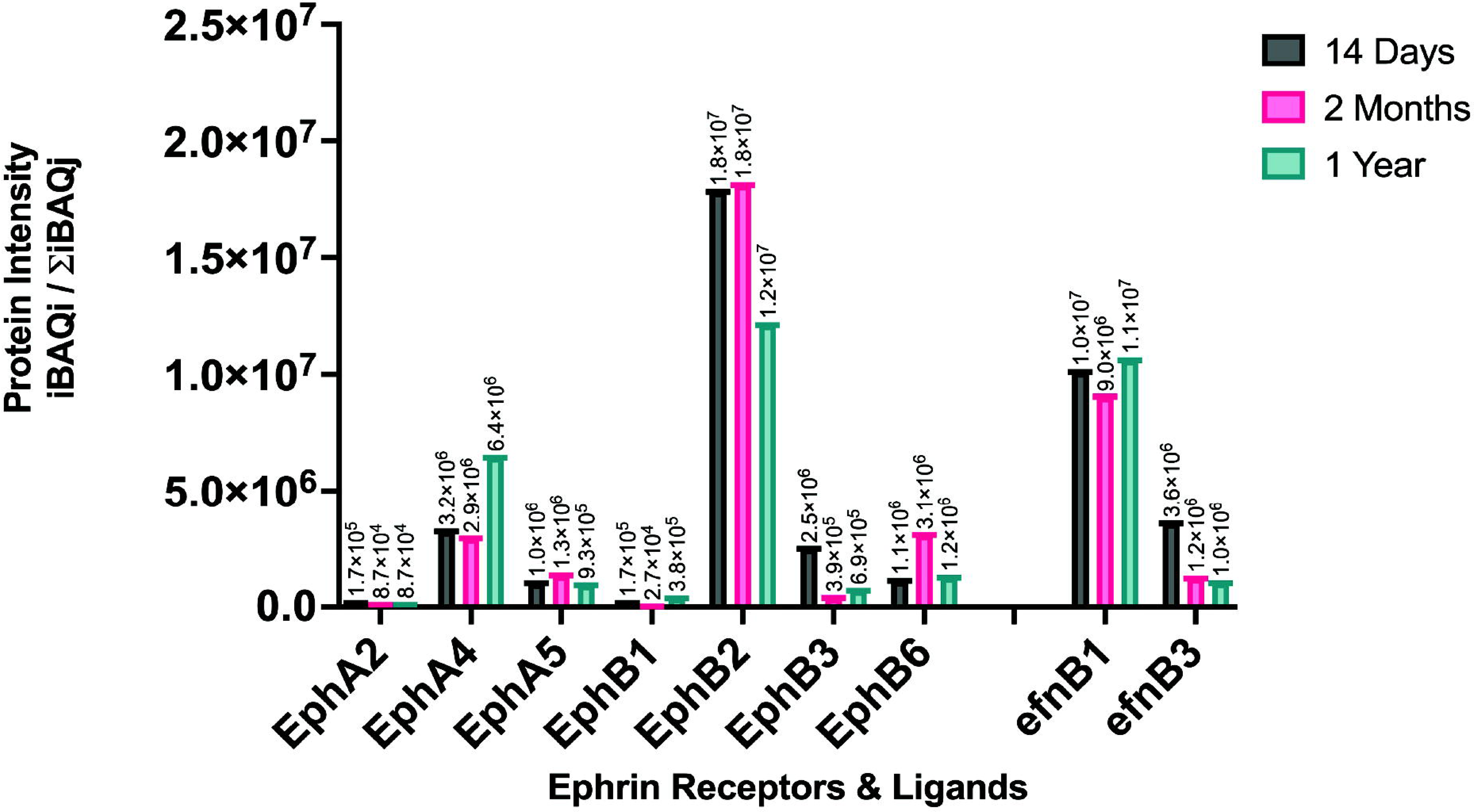
Proteomic analysis of dissected retinal tissue of male C57BL/6J WT mice at 14 days (N = 5 biological replicates, pooled), 2 months (N = 5 biological replicates, pooled), and 12 months of age (N = 5 biological replicates, pooled), using mass spectrometry data independent MS-DIA). Temporal expression of Eph receptors and ephrin ligands in the postnatal retinas.

### Examining Eph Receptor Expression and Activation in Normal and Neuropathic Retinal Tissue

Given the reported literature on ephrin signaling dysregulation in optic neuropathies, we hypothesized that Eph receptor activation would correlate with the onset of neuropathy and precede neuronal loss. To test this hypothesis, we analyzed the proteomic changes in the retina elicited by ONC at 24 and 48hr post-injury when retinal ganglion cell (RGC) loss is not significant in this model [77]. The proteomic analysis conducted on individual Eph receptors revealed substantial alterations in their expression profiles following injury. Specifically, EphA2 exhibited a statistically significant upregulation at the 24-hour time point (*p* = 0.0019), while EphA4 demonstrated a significant increase 48 hours post-injury (*p* = 0.0136). Similarly, EphB1 and EphB3 displayed significant upregulation in expression levels at 24 hours post-injury (p = 0.0213 and p = 0.0005, respectively), while EphB2 exhibited increased expression 48 hours after injury (p = 0.0006) (Supplemental Figure 2A).

Proteomic analysis of phosphorylated Eph receptors unveiled significant increases in the phosphorylation of EphA4 at 24 hours after injury (*p* = 0.0331) and EphA2 at 48 hours after injury (p = 0.0068). EphB1 displayed a significant increase in phosphorylation at 24 hours (*p* = 0.0020) and 48 hours (*p* = 0.0013) after injury. Utilizing a bivalent antibody targeting phosphorylated EphB1+B2, we detected a significant increase in phosphorylation at 24 hours (*p* = 0.0468) and 48 hours (*p* = 0.0039) after injury. Similarly, EphB3 exhibited a significant increase in phosphorylation at both 24 hours (*p* = 0.0228) and 48 hours (*p* = 0.0021) after injury (Supplemental Figure 2B).

The impact of ONC-induced injury on Eph receptor signaling in the retina was investigated through proteomic analysis at 24- and 48-hours post-injury. Eph receptor proteins were characterized and quantified, revealing notable alterations in their phosphorylation levels. Specifically, EphA2 exhibited a significant increase in phosphorylation 48 hours after injury (*p* = 0.0084). Similarly, EphB1 and EphB3 displayed significant increases in phosphorylation at the 48-hour time point following injury (*p* = 0.0053 and *p* = 0.0023, respectively). Additionally, EphA4 demonstrated significant activation both at 24 (*p* = 0.0250) and 48 (*p* < 0.0001) hours after injury (Figure 2).

**Figure 2.**
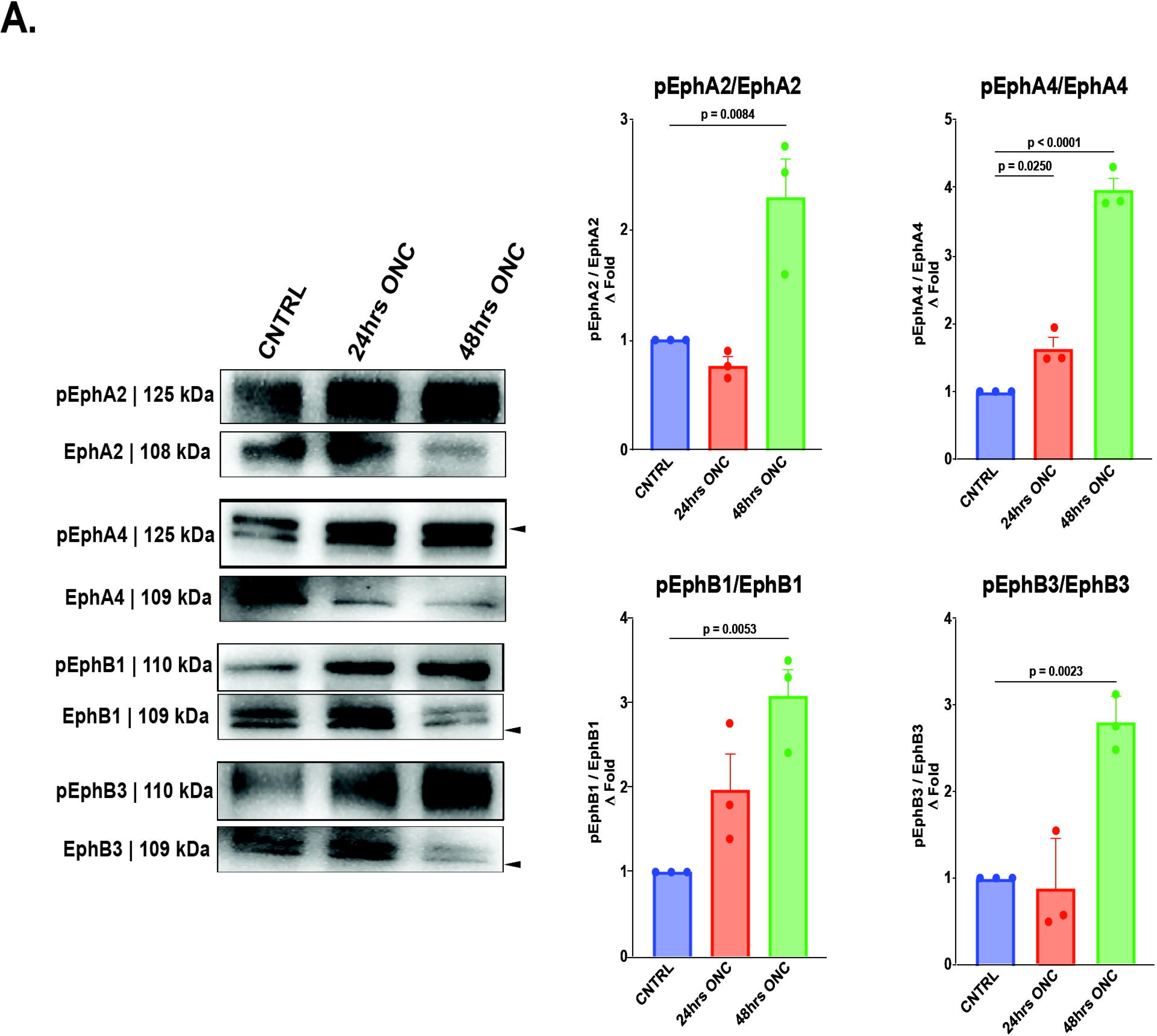
Proteomic quantification of Eph receptors and phosphorylated Eph receptors 24 hours and 48 hours post ONC. (A) Western blot detection and quantification of phosphorylated Eph receptors/ Eph receptors from dissected whole retinal tissue 24 hours and 48 hours post ONC. The geometric means and geometric standard deviations (*N* = 3 biological replicates) are graphed. A one-way ANOVA was applied with a p value of *p* ≤ 0.05 considered statistically significant.

### Distribution of Activated Eph Receptors in the Retinal Layers following ONC and their Association with Neuronal and/or Glial Compartments

Global proteomic analysis and immunoblotting provides insights about the presence of Eph receptors and their relative phosphorylation states respectively, but they do not provide any information about where these receptors reside within the retina. To obtain this information, we used confocal as well as super-resolution microscopy to determine the location and cellular compartmentalization of activated Eph receptors following ONC injury.

Confocal microscopy imaging indicates that phosphorylated EphA2+A3+A4 are distributed exclusively within the inner layers of the retinas (ganglion cell layer to inner plexiform layer) 24 and 48 hours after ONC, and that the fluorescent intensity for these activated receptors is much higher in the injured retina than in non-injured controls at both time points (Figure 3A). Similarly, phosphorylated EphB1+B2 show a similar retinal layer distribution to EphAs and are increased 24 hours and more noticeably 48 hours after ONC when compared to the uninjured retina (Figure 3B).

**Figure 3.**
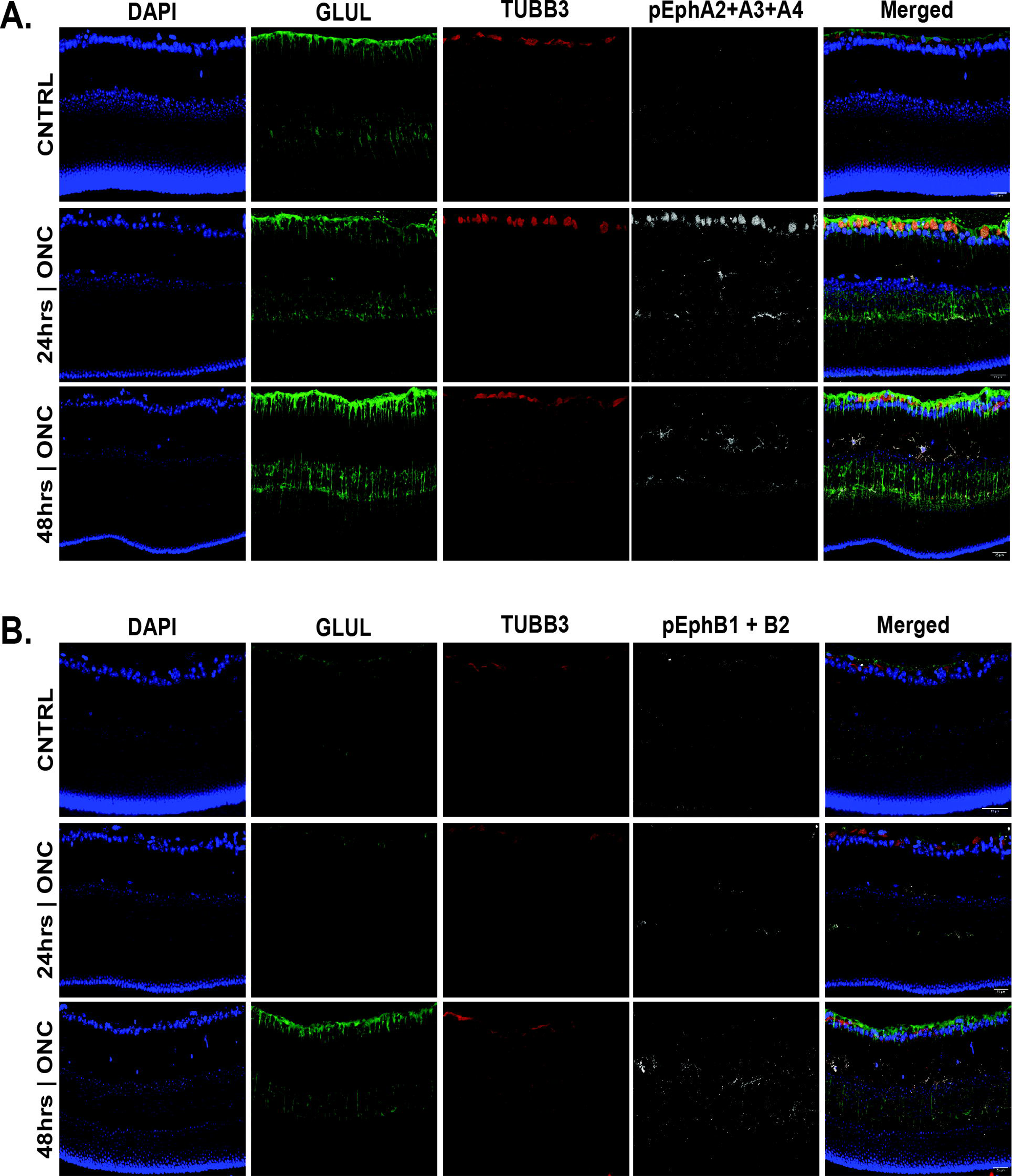
Phosphorylation of multiple EphA- and EphB-class receptors localized within the inner retina in early traumatic optic neuropathy. Immunofluorescent microscopy of retinas (A) Activated Eph receptors A2, A3, and A4, 24 hours and 48 hours post ONC (B) Activated Eph receptors B1 and B2, 24 hours and 48 hours post ONC. Scale bar at 25 μm

STORM imaging followed by OTC co-localization analysis demonstrates that phosphorylated EphA2+A3+A4 at 48 hours post-ONC are significantly more associated to injured neuronal membranes than to uninjured neuronal processes (*p* = 0.003) and significantly more associated to injured neuronal membranes than to injured glial membranes (*p* = 0.0001) (Figure 4 A-C). Similarly, STORM imaging and OTC co-localization analysis show that phosphorylated EphB1+B2 are significantly more co-localized with injured neuronal cells than with uninjured neuronal cells (*p* = 0.0001) and injured neuronal cells than injured glial cells (*p* = 0.0002) (Figure 4 D-F).

**Figure 4.**
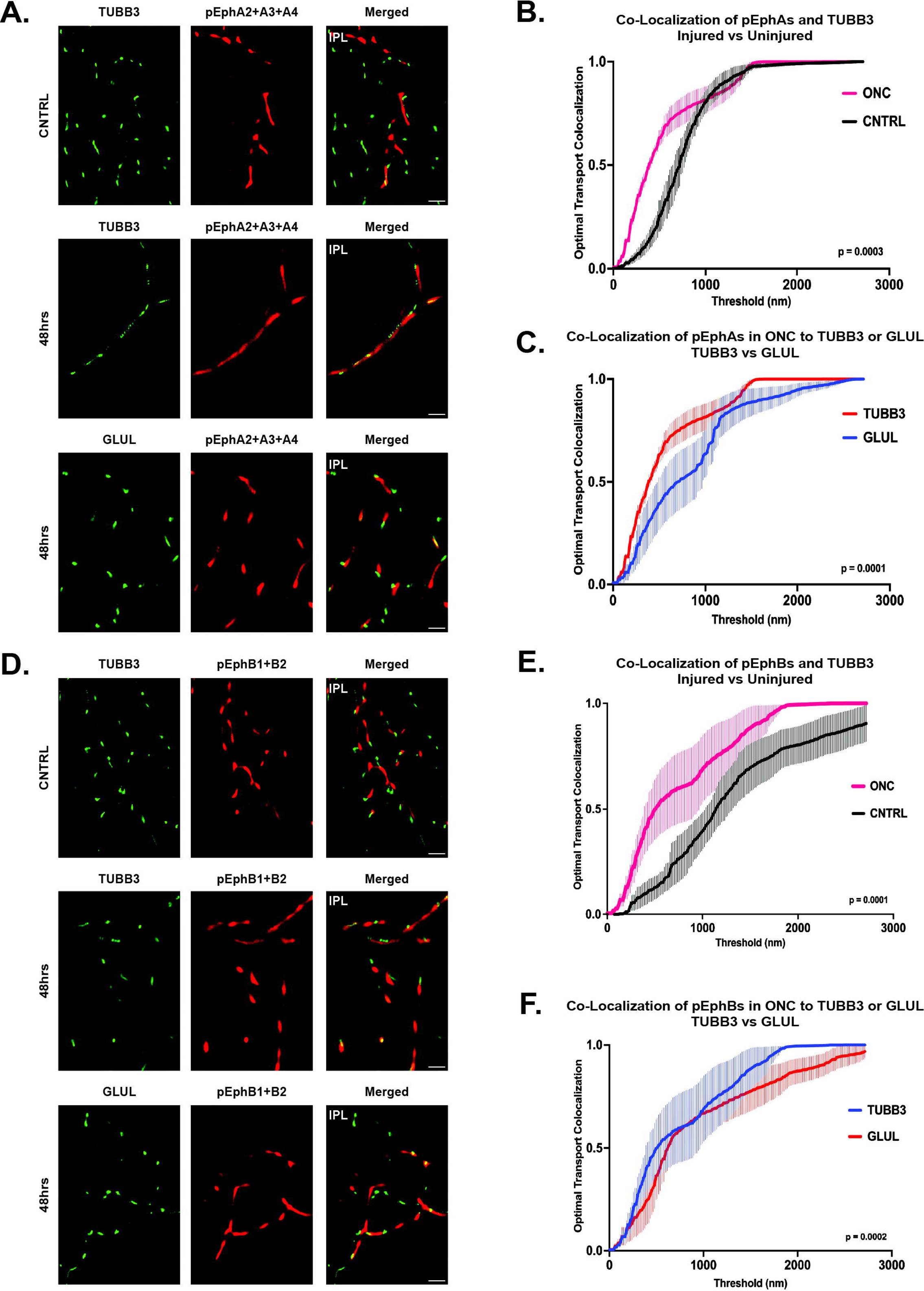
Super-resolution imaging and co-localization analysis of multiple EphA- and EphB-class receptors in neuronal and glial cells within the inner plexiform layer (IPL) of ONC retinas. (A) STORM imaging of phosphorylated Eph receptors A2, A3, and A4 (pEphAs) in 48 hours ONC retinas. (B) OTC analysis comparing the localization of pEphAs to neuronal cells in uninjured and ONC retinas. (C) OTC analysis comparing the localization of pEphAs in glial cells in uninjured vs. ONC retinas. (D) STORM imaging of phosphorylated Eph receptors B1 and B2 (pEphBs) in 48 hours ONC retinas. (B) OTC analysis comparing the localization of pEphBs to neuronal cells in uninjured and ONC retinas. (C) OTC analysis comparing the localization of pEphBs to glial cells in uninjured and ONC retinas. *N* = 3 biological replicates are graphed. Mann-Whitney U tests and a *p* value of *p* ≤ 0.05 is considered statistically significant. Scale bar at 10 μm

### Neuroprotective Effect of Eph Receptor Inhibition

The neuropathic progression is characterized by synaptic instability, retraction, and eventual dendritic and neuronal loss. Our hypothesis suggests that this process may be initiated through the activation of repulsive Eph receptor forward signaling on neuronal membranes. However, the specific contributions of different classes of Eph receptors in this pathological mechanism remain unexplored. Therefore, it is essential to assess which classes may exert a more pronounced role in promoting the neuropathic degeneration of the retina. To investigate this, we propose employing class specific Eph receptor inhibitors to preserve dendritic spines and arbor morphology in the context of neuropathic disease. Evaluating the individual dendritic arbor morphology of retinal ganglion cells (RGCs) necessitates the sparse and selective labeling of individual RGCs, enabling accurate capture and prospective imaging of arbor morphology.

To achieve this, we utilized 2-month-old Thy1-GFP mice, a well-established transgenic mouse model featuring sparsely labeled fluorescence retinal ganglion cells (RGCs) [75,78,79], in our optic nerve crush (ONC) model. We examined the dendritic arbor morphology of Thy1-GFP mice at 48 hours post-injury, a time point when RGC loss is not significant in this model [77], as well as at 6 days post-injury, when significant RGC loss occurs. Our findings indicate that at 48 hours post-injury, there were no significant changes in the dendritic arbor morphology of retinal ganglion cells (*p* = 0.4942), consistent with previous literature [77] (Figure 5A). Strikingly, at 6 days post-injury, a significant decrease in retinal ganglion cell dendritic arbor morphology was observed (*p* = 0.0017), in line with the literature [77] (Figure 5B). These results validate the use of this in vivo model as a platform to evaluate the potential neuroprotective effect of Eph receptor inhibitors.

**Figure 5.**
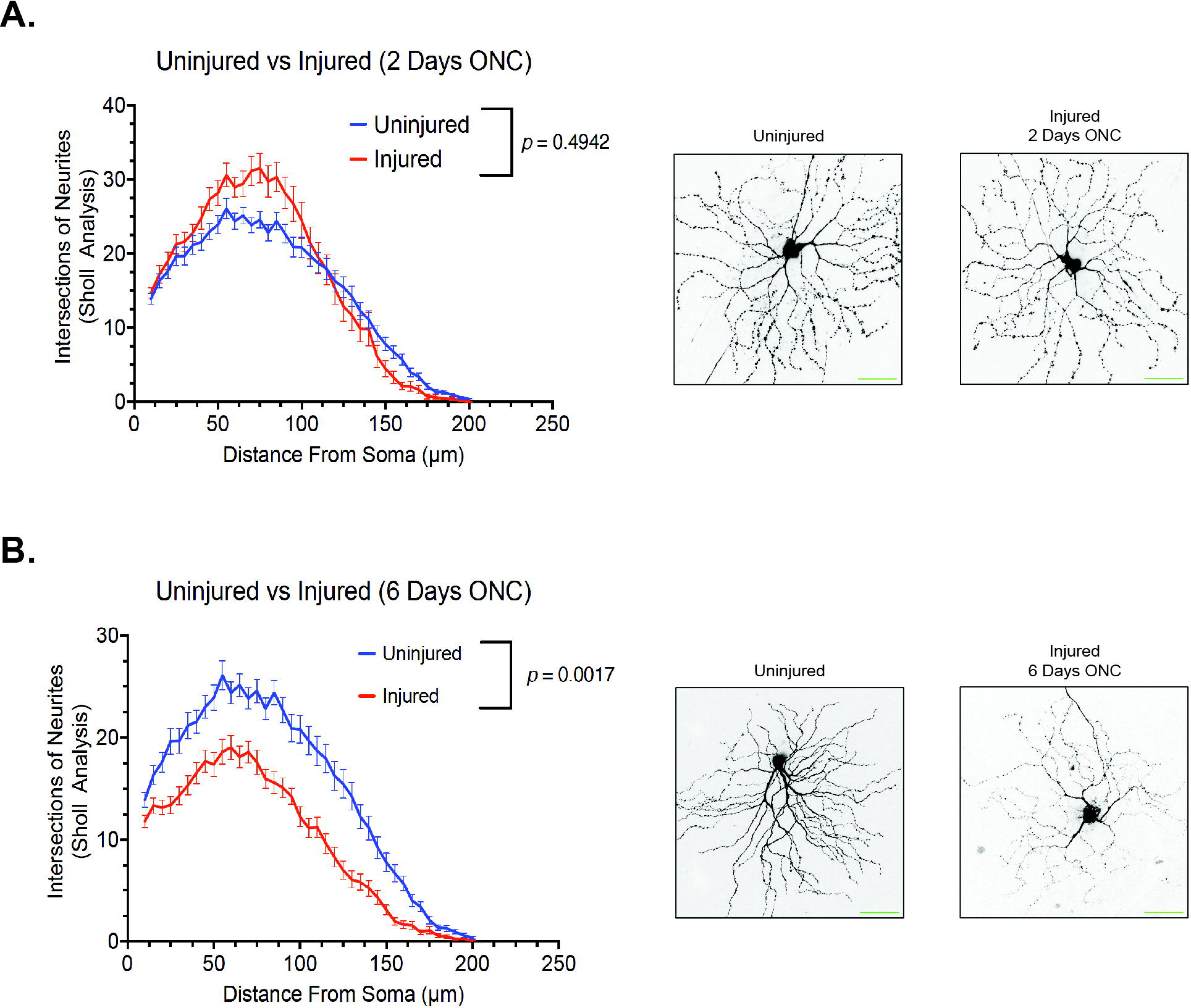
The dynamic change in retinal ganglion cell dendritic arborization detected by confocal microscopy post ONC injury. (A) Quantification of retinal ganglion cell dendritic arborization by sholl analysis 2 days after ONC injury. (*N* = 3 biological replicates. (B) Quantification of retinal ganglion cell dendritic arborization by sholl analysis 6 days after ONC injury. (*N* = 3 biological replicates). Mann-Whitney U tests and a *p* value of *p* ≤ 0.05 is considered statistically significant. Scale bar at 50 μm.

We have shown a correlation between the activation of different Eph receptors and the retraction of neurites and synapses in retinal ganglion cells (RGCs). However, there is currently a lack of pan-Eph receptor inhibitors available in the market. Although specific inhibitors targeting individual Eph receptors do exist, the potential neuroprotective effect of Eph receptor inhibition using commercially available agents, administered intravitreally, remains to be explored. This experiment aimed to evaluate the neuroprotective potential of Eph receptor inhibition on preserving RGC dendritic arborization 6 days after injury. The Eph receptor A inhibitor UniPR129 [80] exhibited a significant neuroprotective effect (*p* = 0.0126) (Figure 6A), while the Eph receptor B inhibitor NVP-BHG712 [81] displayed a greater neuroprotective effect (*p* = 0.0004) (Figure 6B). Strikingly, the most substantial neuroprotection was observed with the combination of both Eph receptor A and B inhibitors (Figure 6C).

**Figure 6.**
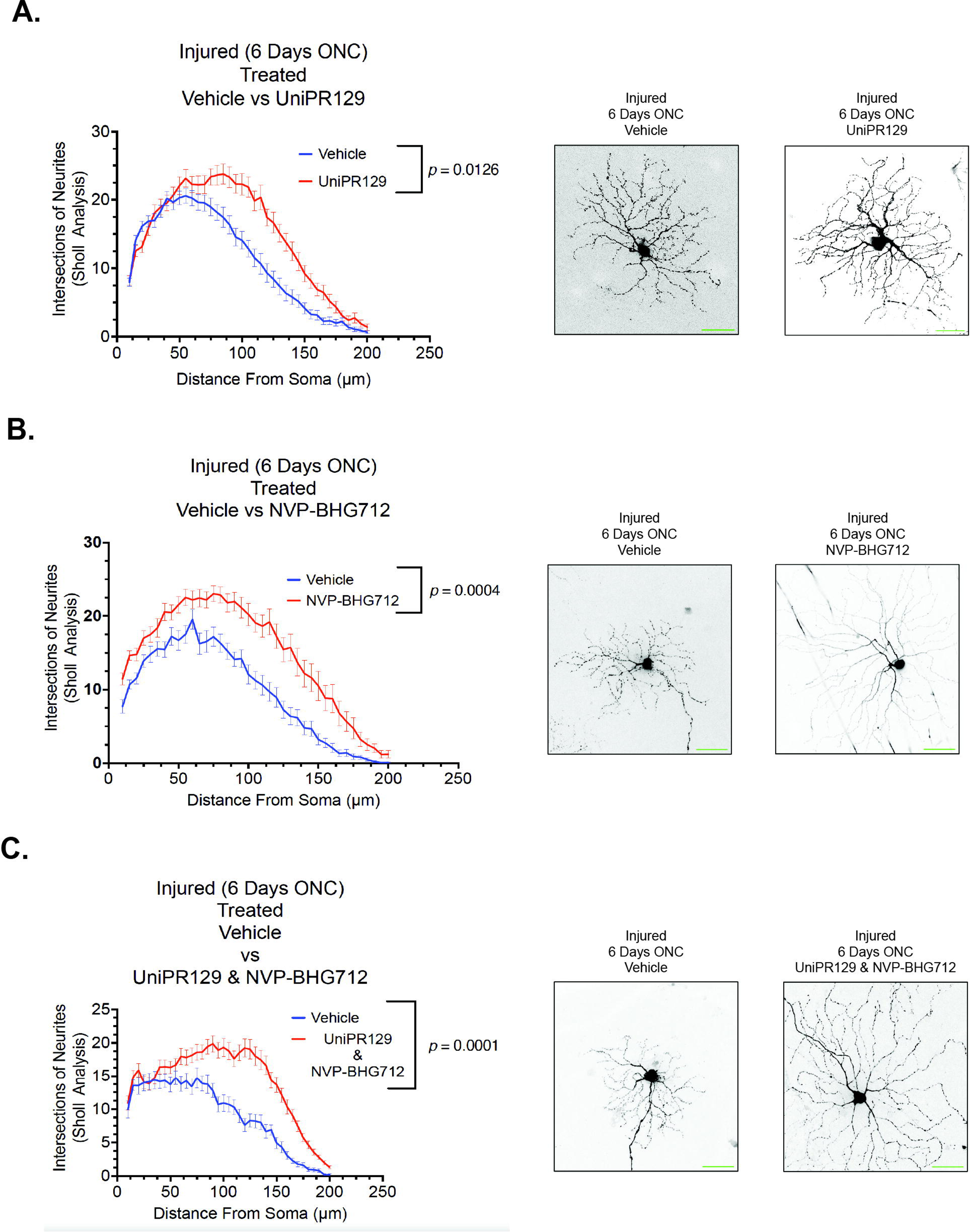
Pan Eph receptor inhibitors demonstrates neuroprotective properties, preserving retinal ganglion cell dendritic arborization as observed by confocal microscopy following ONC injury. (A) Quantification of retinal ganglion cell dendritic arborization by sholl analysis 6 days after ONC injury. Animals treated with 50 μM of UniPR129 (pan Eph receptor A inhibitor), (*N* = 3 biological replicates). (B) Quantification of retinal ganglion cell dendritic arborization by sholl analysis 6 days after ONC injury. Animals treated with 2 μM of NVP- BHG712 (pan Eph receptor B inhibitor), (*N* = 3 biological replicates). (C) Quantification of retinal ganglion cell dendritic arborization by sholl analysis 6 days after ONC injury. Animals treated with a combination of both UniPR129 (50 mM) and NVP-BHG712 (2 mM) pan Eph receptor A&B inhibitors), (*N* = 3 biological replicates). Mann-Whitney U tests and a *p* value of *p* ≤ 0.05 is considered statistically significant. Scale bar at 50 μm.

## DISCUSSION

Central neurodegeneration is a complex and multifactorial process [82–84]. Identifying the molecular determinants underlying the onset and progression of neuropathic states is fundamental to the development of effective treatments. Several lines of evidence show that ‘ephrin signaling’ is one of the most dysregulated pathways in optic neuropathies with varied etiologies [2–5].

Ephrin signaling is highly relevant to the visual system and retina, as counter-gradients of the various members in this ligand-receptor family mediate establish the dorso-ventral and naso-temporal axes during retinotopic map formation in the retina and superior colliculus, as well as guide the decussation of RGC projections through the optic chiasm [11,13,14,47,49–51,85–87]. Ephrin signaling through Eph receptors on RGC projections achieve their guidance by exerting a graded repulsive stimulus on the actin cytoskeletal dynamics across the neuronal membrane [7,31,32,38]. It is remarkable then, that this developmental pathway would be so prominently dysregulated in neuropathic diseases of the visual system. While it can be proposed that the dysregulation constitutes a reactive response by the system in its attempt to re-establish retinotopy, it is interesting to speculate that, given its repulsive and destabilizing nature, activation of the Eph forward signaling pathway may ultimately contribute to neuropathic onset or progression. This study aimed to advance the understanding of dysregulated Eph receptor signaling during early optic neuropathies by addressing key unanswered questions. Firstly, we sought to identify the specific members of this extensive receptor tyrosine kinase (RTK) family that persist in the postnatal retina. Secondly, we aimed to determine which of these receptors are engaged and dysregulated during neuropathic onset. Lastly, we aimed to elucidate the localization of these receptors and their presence on different cellular compartments within the retina. Previous studies have demonstrated the potential regenerative effects in the central nervous system (CNS), particularly the visual system, through modulation of specific Eph receptors [88–90]. Notably, multiple Eph family members have been implicated in the onset and progression of retinal neuropathic diseases (35,38). This is unsurprising considering the involvement of nearly all Eph receptors in the development of retinotopic projections. In light of this, our study aimed to comprehensively explore the contributions of both EphA- and EphB-class receptors using small molecules with established class-specific antagonism: UniPR129 for EphA- class receptors [80] and NVP-BHG712 for EphB-class receptors [81].

Our results show that at least 7 Eph receptors remain patent in the post-natal mouse retina, and that their relative abundance does not change throughout the lifespan of the animals. We further demonstrate that all available Eph receptors become hyperphosphorylated during early time points following optic nerve injury when neuronal dropout has been shown not to be significant. We show that the distribution of Eph receptors is confined to inner retinal structures, from the ganglion cell layer (GCL) to the inner plexiform layer (IPL) and that, within the IPL, activation of Eph receptors occurs on neuronal membranes and not on glial processes upon injury. Finally, we demonstrate that inhibiting ephrin signaling exhibits a significant neuroprotective effect when modulating class specific Eph receptors with the most substantial protection observed with the combination of both Eph receptor A and B inhibitors.

Our study does not allow for the determination of the cause and modality of Eph receptor activation; however, we know that RGCs project their dendritic arbors into the IPL where they synapse with other neurons of the retina. The IPL is also the site for most glial-neuron interactions and synaptic modulation. In Alzheimer’s disease, it has been shown that in the context of neuroinflammation, the approximation of glial-bound ephrin ligands to Eph receptors on synaptic complexes leads to the destabilization and ultimate retraction of the synaptic button [18,19,26,62,91]. As in Alzheimer’s disease, optic neuropathies have a strong neuroinflammatory component to their pathophysiology [92,93], and the loss of synaptic spine density has been shown to precede RGC loss in both glaucoma and traumatic optic neuropathy models [77,94]. Whether glial swelling during optic neuropathy is the cause of Eph receptor activation on RGCs, and whether that constitutes a viable therapeutic target are matters of ongoing investigation by our group. But consistent with this idea, earlier research has demonstrated that ablating an individual Eph receptor has a positive impact on visual function and recovery in optic neuropathy models [89].

In line with our findings, Joly et al. [89] demonstrated that selective deletion of the EphA4 receptor in retinal ganglion cells (RGCs), while leaving the ligand efnA3 unaffected, led to significant regeneration following optic nerve crush. Similarly, Vilallongue et al. [90] reported comparable regenerative effects by knocking down both EphB2 and EphA4, as evidenced by an increase in the number of regenerative events. Our study highlights the aberrant activation of multiple Eph receptors following optic nerve injury, consistent with previous findings reported by Zhou et al. [6], Himanen et al. [95], and Kania et al. [96]. These studies have provided insights into the redundancy and compensatory mechanisms within this highly conserved developmental pathway. Based on these findings, we hypothesized that modulating Eph receptor activity comprehensively or broadly could greatly improve visual outcomes in optic neuropathy. Since there are no currently available pan-Eph receptor inhibitors on the market, we utilized commercially accessible inhibitors for Eph receptor antagonism.

We employed UniPR129 as one of the inhibitors, which is a competitive small molecule that disrupts the binding of EphA-ephrinA [80]. Treatment with UniPR129 demonstrated significant neuroprotection in our optic nerve crush (ONC) injury model (Figure 6A). In addition, we utilized NVP-BHG712, another small molecule that specifically inhibits the EphB kinase domain [81]. Treatment with NVP-BHG712 exhibited even greater neuroprotection in our ONC injury model (Figure 6B). Remarkably, the combined treatment of UniPR129 and NVP-BHG712 (UniPR129 + NVP-BHG712) yielded the most substantial neuroprotection (Figure 6C). These findings strongly support the potential of targeting ephrin signaling as a promising therapeutic strategy for optic neuropathy. Further research is warranted to elucidate the optimal mechanism for modulating ephrin signaling.

The current clinical management of optic neuropathy primarily aims to control disease-associated risk factors, such as elevated intraocular pressure, and slow down disease progression [97–99]. However, advancements in high-throughput assays, coupled with parallel computing and bioinformatics algorithms, now enable the generation of comprehensive system-wide profiles of disease processes and molecular targets. Multiple independent studies have identified dysregulated ‘ephrin signaling’ as a principal component of optic neuropathy pathobiology [3,100]. Remarkably, dysregulation of Eph receptor signaling is detected prior to the onset of visual functional decline, suggesting its involvement in the disease’s pathogenesis rather than being merely another risk factor. These observations hold significant importance and require further exploration to understand their role in neuropathic progression. Identifying appropriate molecular targets is crucial for developing effective therapies for these conditions. By focusing on active ephrin forward signaling as a molecular determinant of neuropathic progression in the visual system, we aim to establish a framework for novel treatments that preserve and restore vision in patients with these debilitating conditions.

## CONCLUSIONS

This study demonstrates that all detectable Eph receptors within the postnatal murine retina become aberrantly hyperactivated on neuronal membranes within the inner plexiform layer in the acute phase of optic neuropathic onset, and prior to a significant decline in retinal ganglion cell (RGC) numbers following optic nerve crush injury. Given the strong repulsive and destabilizing effect that Eph forward signaling exerts on neuronal processes, these results constitute a significant advance in our characterization of the molecular determinants of neuropathic diseases of the visual system and underscores the need to further elucidate the role that Eph receptor signaling plays in disease progression and its value as a therapeutic target.

## LIST OF ABBREVIATIONS

CNS: Central Nervous System; efn: ephrin ligands; Eph: Eph receptors; GCL: ganglion cell layer; IPL: inner plexiform layer; MS-DIA: Mass Spectrometry Data Independent Acquisition; ONC: Optic Nerve Crush; ON: Optic Neuropathy; OTC: Optical Transport Colocalization; RGC: retinal ganglion cell; receptor tyrosine kinase (RTK); STORM: Stochastic Optical Reconstruction Microscopy; TBST: Tris-buffered saline with 0.1% Tween 20

## DECLARATIONS SECTION

### Ethical Approval and Consent to Participate

Animals were treated in accordance with the National Research Council’s Guide for the Care and Use of Laboratory Animals and the ARVO Statement for the Use of Animals in Ophthalmic and Vision Research. Animal procedures were approved by the Institutional Animal Care and Use Committee at the University of Miami.

### Consent For Publication

Not Applicable

### Availability Of Data and Materials

The dataset(s) supporting the conclusions of this article is(are) available in the Mendeley Data repository, Strong, Tom (2023), “Strong 2023 Eph Receptors”, Mendeley Data, V1, doi: 10.17632/zcpzvjb2vw.1

## Funding

This work was supported in part by a generous philanthropic gift from Dr. Nasser Ibrahim Al-Rashid to the Bascom Palmer Eye Institute, and an Alcon Research Institute Young Investigator Grant (DP). The Bascom Palmer Eye Institute is supported by NIH Center Core Grant P30EY01801 and a Research to Prevent Blindness Unrestricted Grant (New York, NY, USA).

### Authors’ Contributions

TAS designed and conducted the animal and molecular experiments and was responsible for data collection, analysis, and interpretation. TAS and DP wrote the manuscript. JE and QW assisted in data collection, analysis, and interpretation. HW assisted in the establishment of all animal experiments. PL and GG assisted in data collection and analysis. DP and DT coordinated and directed the entire project. All authors read and approved the final manuscript.

## Acknowledgments

Figures were created in part with Biorender.com. We would like to thank Dr. Galina Dvoriantchikova for her assistance in coordinating animal experimentations. We thank Dr. Ramin Shiekhattar for his assistance with STORM super resolution imaging.

---

**Supplementary Figure 1.**
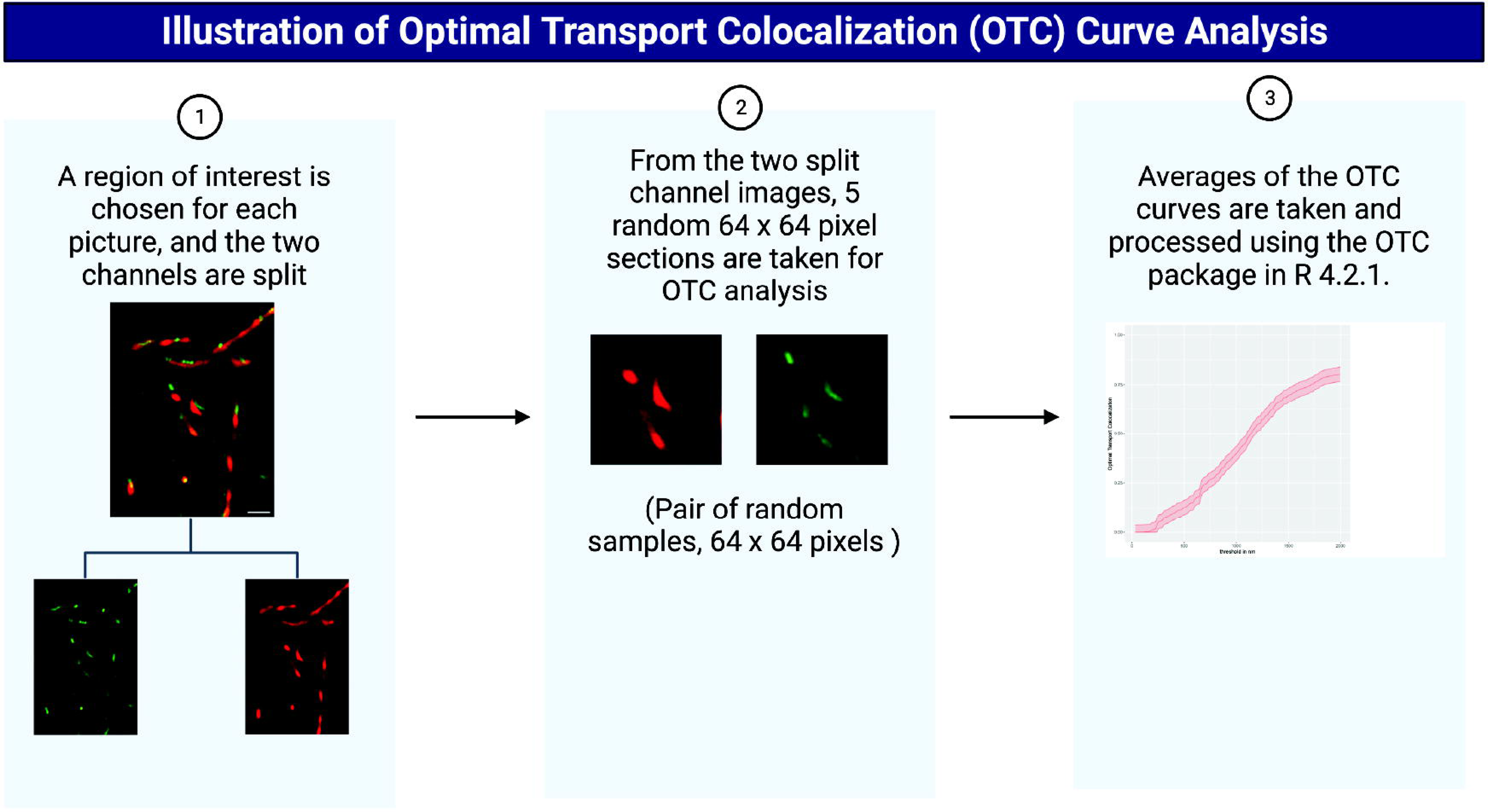
Illustration of Optimal Transport Colocalization curve analysis workflow. Created with BioRender.com

**Supplementary Figure 2.**
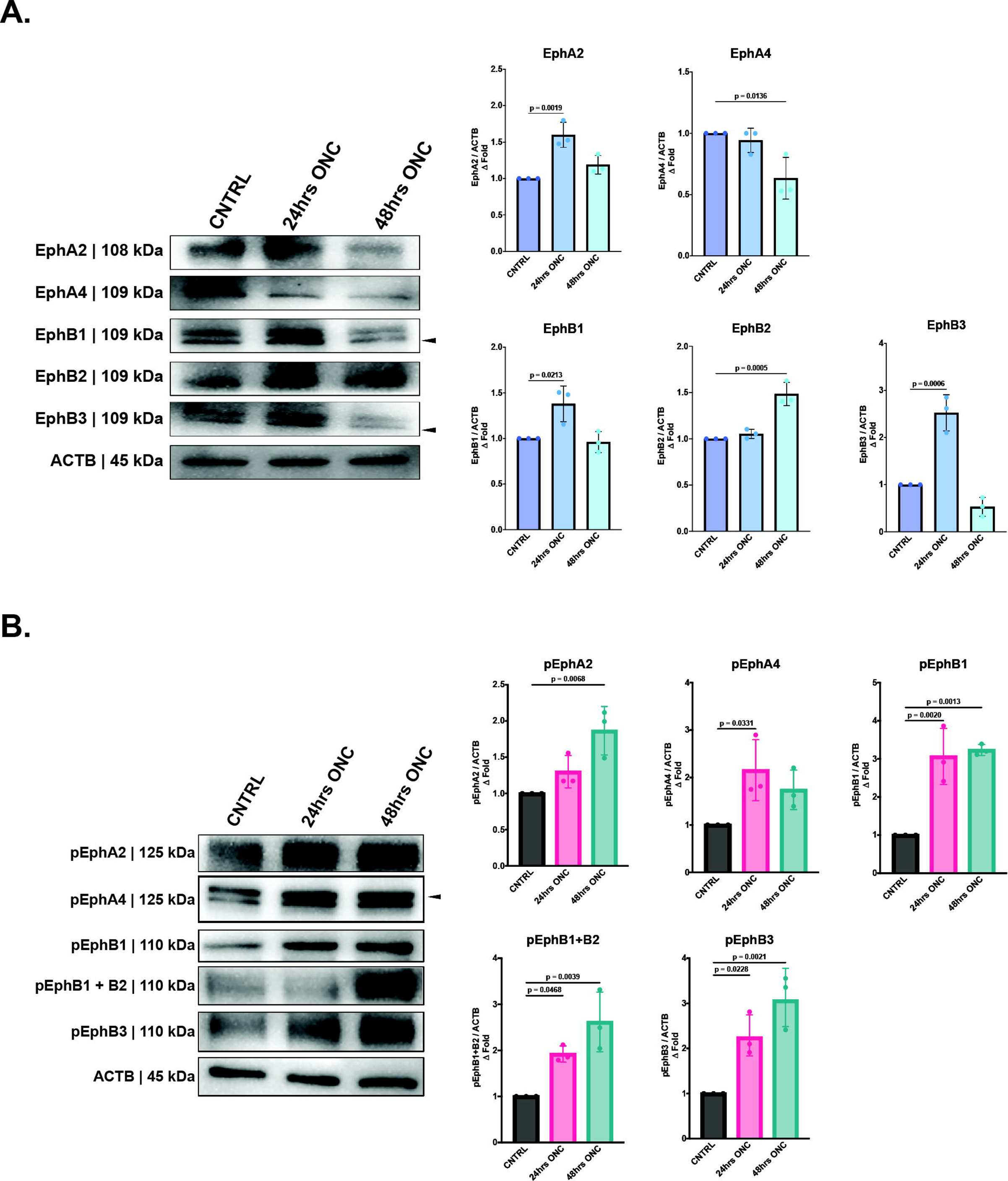
Proteomic quantification of Eph receptors and phosphorylated Eph receptors 24 hours and 48 hours post ONC. (A) Western blot detection and quantification of phosphorylated Eph receptors/b-actin from dissected whole retinal tissue 24 hours and 48 hours post ONC. The geometric means and geometric standard deviations (*N* = 3 biological replicates) are graphed. (B) Western blot detection and quantification of Eph receptors/b-actin from dissected whole retinal tissue 24 hours and 48 hours post ONC. The geometric means and geometric standard deviations (N = 3 biological replicates) are graphed.

**Supplemental Table 1.**
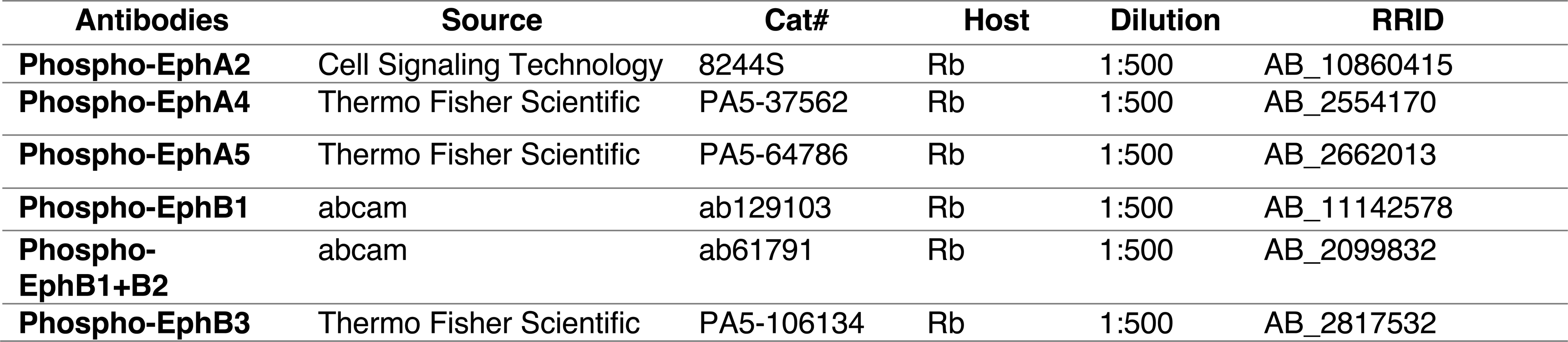
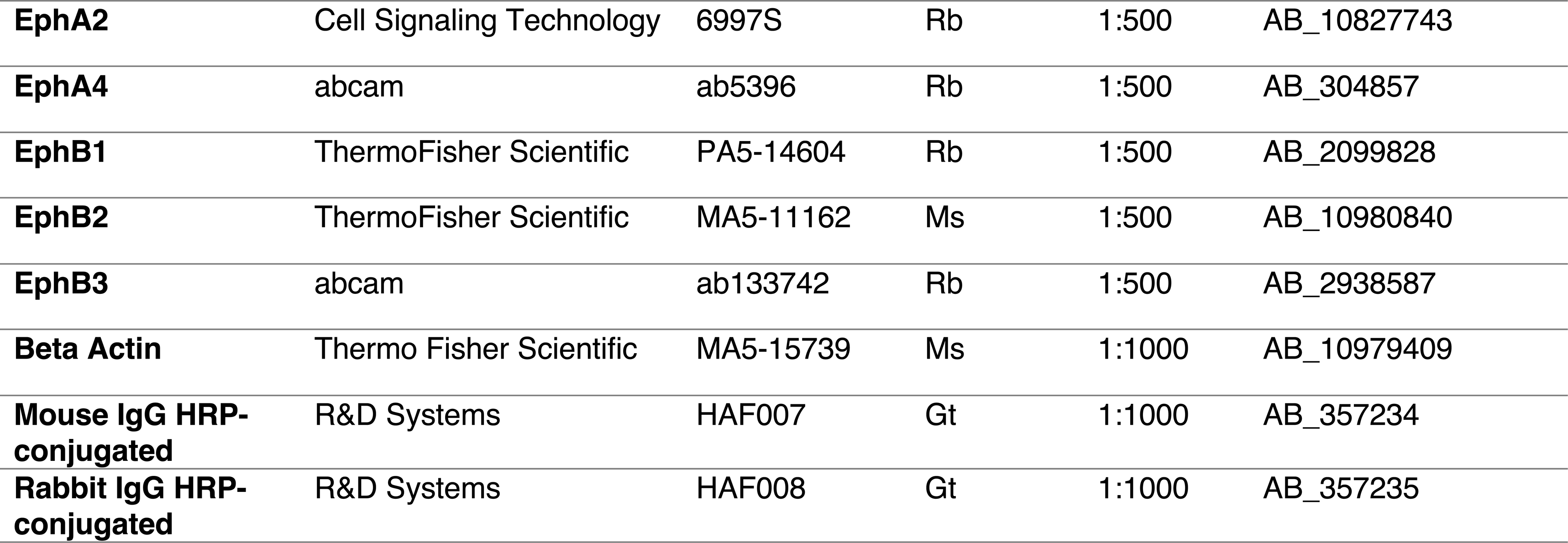
Antibodies Used for Western Blotting

| Antibodies | Source | Cat# | Host | Dilution | RRID |
| --- | --- | --- | --- | --- | --- |
| **Phospho-EphA2** | Cell Signaling Technology | 8244S | Rb | 1:500 | AB_10860415 |
| **Phospho-EphA4** | Thermo Fisher Scientific | PA5-37562 | Rb | 1:500 | AB_2554170 |
| **Phospho-EphA5** | Thermo Fisher Scientific | PA5-64786 | Rb | 1:500 | AB_2662013 |
| **Phospho-EphB1** | abcam | ab129103 | Rb | 1:500 | AB_11142578 |
| **Phospho-EphB1+B2** | abcam | ab61791 | Rb | 1:500 | AB_2099832 |
| **Phospho-EphB3** | Thermo Fisher Scientific | PA5-106134 | Rb | 1:500 | AB_2817532 |
| **EphA2** | Cell Signaling Technology | 6997S | Rb | 1:500 | AB_10827743 |
| **EphA4** | abcam | ab5396 | Rb | 1:500 | AB_304857 |
| **EphB1** | ThermoFisher Scientific | PA5-14604 | Rb | 1:500 | AB_2099828 |
| **EphB2** | ThermoFisher Scientific | MA5-11162 | Ms | 1:500 | AB_10980840 |
| **EphB3** | abcam | ab133742 | Rb | 1:500 | AB_2938587 |
| **Beta Actin** | Thermo Fisher Scientific | MA5-15739 | Ms | 1:1000 | AB_10979409 |
| **Mouse IgG HRP-conjugated** | R&D Systems | HAF007 | Gt | 1:1000 | AB_357234 |
| **Rabbit IgG HRP-conjugated** | R&D Systems | HAF008 | Gt | 1:1000 | AB_357235 |

**Supplemental Table 2.** Antibodies Used for Immunofluorescence and STORM Staining

| **Antibodies** | **Source** | **Cat#** | **Host** | **Dilution** | **RRID** |
| --- | --- | --- | --- | --- | --- |
| **Phospho-EphA2+A3+A4** | abcam | ab62256 | Rb | 1:100 | AB_942240 |
| **Phospho-EphB1+B2** | abcam | ab61791 | Rb | 1:100 | AB_2099832 |
| **Anti-beta III Tubulin** | Novus Biologicals | NB100-1612 | Ck | 1:100 | AB_10000548 |
| **Anti-Glutamine Synthetase** | Millipore Sigma | MAB302 | Ms | 1:100 | AB_2110656 |
| **Alexa Fluor 488<sup>IF</sup>** | abcam | ab150113 | Gt | 1:100 | AB_2576208 |
| **Alexa Fluor 555<sup>IF</sup>** | abcam | ab150170 | Gt | 1:100 | AB_2893330 |
| **Alexa Fluor 647<sup>IF</sup>** | abcam | ab150079 | Gt | 1:100 | AB_2722623 |
| **Janelia Fluor 646<sup>STORM</sup>** | Novus Biologicals | NBP1-72732JF646 | Gt | 1:100 | AB_2936900 |
| **Alexa Fluor 568<sup>STORM</sup>** | ThermoFisher Scientific | A-11004 | Gt | 1:100 | AB_2534072 |

**Supplemental Table 3.**
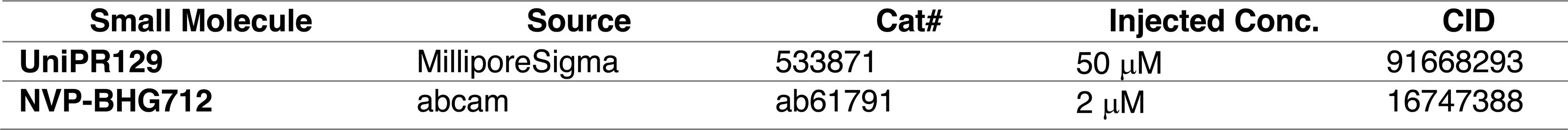
Eph Receptor Inhibitors Used in Intravitreal injections

| **Small Molecule** | **Source** | **Cat#** | **Injected Conc.** | **CID** |
| --- | --- | --- | --- | --- |
| **UniPR129** | MilliporeSigma | 533871 | 50 μM | 91668293 |
| **NVP-BHG712** | abcam | ab61791 | 2 μM | 16747388 |

## REFERENCE

1. Health DYCE, 2003. Retinal diseases and vision 2020. ncbi.nlm.nih.gov.

2. Colak D, Morales J, Bosley TM, Al-Bakheet A, AlYounes B, Kaya N, et al. Genome-Wide Expression Profiling of Patients with Primary Open Angle GlaucomaGene Expression Profiling of POAG. Investigative Ophthalmology & Visual Science. 2012;53:5899–6.

3. Nikolskaya T, Nikolsky Y, Serebryiskaya T, Zvereva S, Sviridov E, Dezso Z, et al. Network analysis of human glaucomatous optic nerve head astrocytes. BMC Medical Genomics [Internet]. 2009;2:389–26. Available from: https://bmcmedgenomics.biomedcentral.com/track/pdf/10.1186/1755-8794-2-24

4. Williams PA, Harder JM, Foxworth NE, Cardozo BH, Cochran KE, John SWM. Nicotinamide and WLDS Act Together to Prevent Neurodegeneration in Glaucoma. Front Neurosci-switz. 2017;11:232.

5. Tezel G, Thornton IL, Tong MG, Luo C, Yang X, Cai J, et al. Immunoproteomic Analysis of Potential Serum Biomarker Candidates in Human Glaucoma. Investigative Ophthalmology & Visual Science [Internet]. 2012;53:8222–10. Available from: https://www.ncbi.nlm.nih.gov/pmc/articles/PMC3522442/pdf/i1552-5783-53-13-8222.pdf

6. Zhou R. The Eph Family Receptors and Ligands. Pharmacol Therapeut. 1998;77:151– 81.

7. Orioli D, Klein R. The eph receptor family:axonal guidance by contact repulsion. Trends Genet. 1997;13:354–9.

8. Mellott DO, Burke RD. The molecular phylogeny of eph receptors and ephrin ligands. Bmc Cell Biol. 2008;9:27.

9. Murai KK, Pasquale EB. Eph Receptors, Ephrins, and Synaptic Function. Neurosci. 2004;10:304–14.

10. Wu Z, Ashlin TG, Xu Q, Wilkinson DG. Role of forward and reverse signaling in Eph receptor and ephrin mediated cell segregation. Exp Cell Res. 2019;381:57–65.

11. Brown A, Yates PA, Burrola P, Ortuño D, Vaidya A, Jessell TM, et al. Topographic Mapping from the Retina to the Midbrain Is Controlled by Relative but Not Absolute Levels of EphA Receptor Signaling. Cell. 2000;102:77–88.

12. Feldheim DA, O’Leary DDM. Visual Map Development: Bidirectional Signaling, Bifunctional Guidance Molecules, and Competition. Csh Perspect Biol. 2010;2:a001768.

13. Davenport RW, Thies E, Zhou R, Nelson PG. Cellular Localization of Ephrin-A2, Ephrin-A5, and Other Functional Guidance Cues Underlies Retinotopic Development across Species. J Neurosci. 1998;18:975–86.

14. Tessier-Lavigne M. Eph receptor tyrosine kinases, axon repulsion, and the development of topographic maps. Cell. 1995;82:345–8.

15. Cang J, Kaneko M, Yamada J, Woods G, Stryker MP, Feldheim DA. Ephrin-As Guide the Formation of Functional Maps in the Visual Cortex. Neuron. 2005;48:577–89.

16. Feldheim DA, Vanderhaeghen P, Hansen MJ, Frisén J, Lu Q, Barbacid M, et al. Topographic Guidance Labels in a Sensory Projection to the Forebrain. Neuron. 1998;21:1303–13.

17. Zhou L, Jones EV, Murai KK. EphA Signaling Promotes Actin-based Dendritic Spine Remodeling through Slingshot Phosphatase. Journal of Biological Chemistry [Internet]. 2012;287:9346–59. Available from: https://www.jbc.org/content/287/12/9346.full.pdf

18. Carmona MA, Murai KK, Wang L, Roberts AJ, Pasquale EB. Glial ephrin-A3 regulates hippocampal dendritic spine morphology and glutamate transport. Proc National Acad Sci. 2009;106:12524–9.

19. Murai KK, Nguyen LN, Irie F, Yamaguchi Y, Pasquale EB. Control of hippocampal dendritic spine morphology through ephrin-A3/EphA4 signaling. Nat Neurosci. 2003;6:153–60.

20. Bourgin C, Murai KK, Richter M, Pasquale EB. The EphA4 receptor regulates dendritic spine remodeling by affecting β1-integrin signaling pathways. J Cell Biology. 2007;178:1295–307.

21. Chen Y, Fu AKY, Ip NY. Eph receptors at synapses: Implications in neurodegenerative diseases. Cell Signal. 2012;24:606–11.

22. Irie F, Yamaguchi Y. EPHB receptor signaling in dendritic spine development. Front Biosci. 2004;9:1365.

23. Penzes P, Beeser A, Chernoff J, Schiller MR, Eipper BA, Mains RE, et al. Rapid Induction of Dendritic Spine Morphogenesis by trans-Synaptic EphrinB-EphB Receptor Activation of the Rho-GEF Kalirin. Neuron. 2003;37:263–74.

24. Thompson SM. Ephrins keep dendritic spines in shape. Nat Neurosci. 2003;6:103–4.

25. Grunwald IC, Korte M, Wolfer D, Wilkinson GA, Unsicker K, Lipp H-P, et al. Kinase-Independent Requirement of EphB2 Receptors in Hippocampal Synaptic Plasticity. Neuron. 2001;32:1027–40.

26. Murai KK, Pasquale EB. Eph receptors and ephrins in neuron–astrocyte communication at synapses. Glia. 2011;59:1567–78.

27. Coonan JR, Greferath U, Messenger J, Hartley L, Murphy M, Boyd AW, et al. Development and reorganization of corticospinal projections in EphA4 deficient mice. J Comp Neurol. 2001;436:248–62.

28. Fei E, Xiong W-C, Mei L. Ephrin-B3 recruits PSD-95 to synapses. Nat Neurosci. 2015;18:1535–7.

29. Hruska M, Henderson NT, Xia NL, Marchand SJL, Dalva MB. Anchoring and synaptic stability of PSD-95 is driven by ephrin-B3. Nat Neurosci. 2015;18:1594–605.

30. Marquardt T, Shirasaki R, Ghosh S, Andrews SE, Carter N, Hunter T, et al. Coexpressed EphA Receptors and Ephrin-A Ligands Mediate Opposing Actions on Growth Cone Navigation from Distinct Membrane Domains. Cell. 2005;121:127–39.

31. Hansen MJ, Dallal GE, Flanagan JG. Retinal Axon Response to Ephrin-As Shows a Graded, Concentration-Dependent Transition from Growth Promotion to Inhibition. Neuron. 2004;42:717–30.

32. Birgbauer E, Oster SF, Severin CG, Sretavan DW. Retinal axon growth cones respond to EphB extracellular domains as inhibitory axon guidance cues. Development. 2001;128:3041–8.

33. Wahl S, Barth H, Ciossek T, Aktories K, Mueller BK. Ephrin-A5 Induces Collapse of Growth Cones by Activating Rho and Rho Kinase. J Cell Biology. 2000;149:263–70.

34. Sahin M, Greer PL, Lin MZ, Poucher H, Eberhart J, Schmidt S, et al. Eph-Dependent Tyrosine Phosphorylation of Ephexin1 Modulates Growth Cone Collapse. Neuron. 2005;46:191–204.

35. Cox EC, Müller B, Bonhoeffer F. Axonal guidance in the chick visual system: Posterior tectal membanes induce collapse of growth cones from the temporal retina. Neuron. 1990;4:31–7.

36. Shamah SM, Lin MZ, Goldberg JL, Estrach S, Sahin M, Hu L, et al. EphA Receptors Regulate Growth Cone Dynamics through the Novel Guanine Nucleotide Exchange Factor Ephexin. Cell. 2001;105:233–44.

37. Philipsborn AC von, Lang S, Loeschinger J, Bernard A, David C, Lehnert D, et al. Growth cone navigation in substrate-bound ephrin gradients. Development. 2006;133:2487–95.

38. Mendes SW, Henkemeyer M, Liebl DJ. Multiple Eph Receptors and B-Class Ephrins Regulate Midline Crossing of Corpus Callosum Fibers in the Developing Mouse Forebrain. J Neurosci. 2006;26:882–92.

39. Chenaux G, Henkemeyer M. Forward signaling by EphB1/EphB2 interacting with ephrin-B ligands at the optic chiasm is required to form the ipsilateral projection. Eur J Neurosci. 2011;34:1620–33.

40. Wang L-C, Rachel RA, Marcus RC, Mason CA. Chemosuppression of Retinal Axon Growth by the Mouse Optic Chiasm. Neuron. 1996;17:849–62.

41. Wizenmann A, Thanos S, Boxberg Y von, Bonhoeffer F. Differential reaction of crossing and non-crossing rat retinal axons on cell membrane preparations from the chiasm midline: an in vitro study. Dev Camb Engl. 1993;117:725–35.

42. Williams SE, Mann F, Erskine L, Sakurai T, Wei S, Rossi DJ, et al. Ephrin-B2 and EphB1 Mediate Retinal Axon Divergence at the Optic Chiasm. Neuron [Internet]. 2003;39:919–35. Available from: https://ac.els-cdn.com/S0896627303005415/1-s2.0- S0896627303005415-main.pdf?_tid=2248fb2b-6503-4f6c-b509-585f5fcabf8c&acdnat=1549052090_71e40e9478e0bd96bc647c33db92a5e8

43. Fu W-Y, Chen Y, Sahin M, Zhao X-S, Shi L, Bikoff JB, et al. Cdk5 regulates EphA4- mediated dendritic spine retraction through an ephexin1-dependent mechanism. Nat Neurosci. 2007;10:67–76.

44. Vargas LM, Leal N, Estrada LD, González A, Serrano F, Araya K, et al. EphA4 Activation of c-Abl Mediates Synaptic Loss and LTP Blockade Caused by Amyloid-β Oligomers. Plos One. 2014;9:e92309.

45. Naj AC, Jun G, Beecham GW, Wang L-S, Vardarajan BN, Buros J, et al. Common variants at MS4A4/MS4A6E, CD2AP, CD33 and EPHA1 are associated with late-onset Alzheimer’s disease. Nat Genet. 2011;43:436–41.

46. Perez EJ, Cepero ML, Perez SU, Coyle JT, Sick TJ, Liebl DJ. EphB3 signaling propagates synaptic dysfunction in the traumatic injured brain. Neurobiology of Disease. 2016;94:73–84.

47. Braisted JE, McLaughlin T, Wang HU, Friedman GC, Anderson DJ, O’leary DDM. Graded and Lamina-Specific Distributions of Ligands of EphB Receptor Tyrosine Kinases in the Developing Retinotectal System. Dev Biol. 1997;191:14–28.

48. Frisén J, Yates PA, McLaughlin T, Friedman GC, O’Leary DDM, Barbacid M. Ephrin- A5 (AL-1/RAGS) Is Essential for Proper Retinal Axon Guidance and Topographic Mapping in the Mammalian Visual System. Neuron. 1998;20:235–43.

49. O’Leary DDM, McLaughlin T. Mechanisms of retinotopic map development: Ephs, ephrins, and spontaneous correlated retinal activity. Prog Brain Res. 2005;147:43–65.

50. Liu M, Wang L, Cang J. Different roles of axon guidance cues and patterned spontaneous activity in establishing receptive fields in the mouse superior colliculus. Front Neural Circuit. 2014;8:23.

51. Wilkinson DG. Topographic mapping: Organising by repulsion and competition? Curr Biol. 2000;10:R447–51.

52. Fu AKY, Hung K-W, Huang H, Gu S, Shen Y, Cheng EYL, et al. Blockade of EphA4 signaling ameliorates hippocampal synaptic dysfunctions in mouse models of Alzheimer’s disease. Proc National Acad Sci. 2014;111:9959–64.

53. Suzuki K, Aimi T, Ishihara T, Mizushima T. Identification of approved drugs that inhibit the binding of amyloid β oligomers to ephrin type-B receptor 2. Febs Open Bio. 2016;6:461–8.

54. Inoue E, Deguchi-Tawarada M, Togawa A, Matsui C, Arita K, Katahira-Tayama S, et al. Synaptic activity prompts γ-secretase–mediated cleavage of EphA4 and dendritic spine formation. J Cell Biology. 2009;185:551–64.

55. Morgan K. The three new pathways leading to Alzheimer’s disease. Neuropath Appl Neuro. 2011;37:353–7.

56. Du J, Tran T, Fu C, Sretavan DW. Upregulation of EphB2 and ephrin-B2 at the Optic Nerve Head of DBA/2J Glaucomatous Mice Coincides with Axon Loss. Invest Ophth Vis Sci. 2007;48:5567–81.

57. Dong L-D, Gao F, Wang X-H, Miao Y, Wang S-Y, Wu Y, et al. GluA2 Trafficking Is Involved in Apoptosis of Retinal Ganglion Cells Induced by Activation of EphB/EphrinB Reverse Signaling in a Rat Chronic Ocular Hypertension Model. J Neurosci. 2015;35:5409–21.

58. Fu CT, Tran T, Sretavan D. Axonal/Glial Upregulation of EphB/ephrin-B Signaling in Mouse Experimental Ocular Hypertension. Invest Ophth Vis Sci. 2010;51:991–1001.

59. Liu S-T, Zhong S-M, Li X-Y, Gao F, Li F, Zhang M-L, et al. EphrinB/EphB forward signaling in Müller cells causes apoptosis of retinal ganglion cells by increasing tumor necrosis factor alpha production in rat experimental glaucomatous model. Acta Neuropathologica Communications [Internet]. 2018;6:1–15. Available from: https://www.ncbi.nlm.nih.gov/pmc/articles/PMC6201539/pdf/40478_2018_Article_618.pdf

60. Lukas TJ, Miao H, Chen L, Riordan SM, Li W, Crabb AM, et al. Susceptibility to glaucoma: differential comparison of the astrocyte transcriptome from glaucomatous African American and Caucasian American donors. Genome Biol. 2008;9:R111.

61. Tezel G. A proteomics view of the molecular mechanisms and biomarkers of glaucomatous neurodegeneration. Prog Retin Eye Res. 2013;35:18–43.

62. Nikolakopoulou AM, Koeppen J, Garcia M, Leish J, Obenaus A, Ethell IM. Astrocytic Ephrin-B1 Regulates Synapse Remodeling Following Traumatic Brain Injury. ASN Neuro [Internet]. 2016;8:175909141663022–18. Available from: https://www.ncbi.nlm.nih.gov/pmc/articles/PMC4774052/pdf/10.1177_17590914166302 20.pdf

63. Frugier T, Conquest A, McLean C, Currie P, Moses D, Goldshmit Y. Expression and Activation of EphA4 in the Human Brain After Traumatic Injury. J Neuropathology Exp Neurology. 2012;71:242–50.

64. Assis-Nascimento P, Tsenkina Y, Liebl DJ. EphB3 signaling induces cortical endothelial cell death and disrupts the blood–brain barrier after traumatic brain injury. Cell Death Dis. 2018;9:7.

65. Ernst A-S, Böhler L-I, Hagenston AM, Hoffmann A, Heiland S, Sticht C, et al. EphB2- dependent signaling promotes neuronal excitotoxicity and inflammation in the acute phase of ischemic stroke. Acta Neuropathologica Commun. 2019;7:15.

66. Lemmens R, Jaspers T, Robberecht W, Thijs VN. Modifying expression of EphA4 and its downstream targets improves functional recovery after stroke. Hum Mol Genet. 2013;22:2214–20.

67. Chen F, Liu Z, Peng W, Gao Z, Cao Z, Yan T, et al. Activation of EphA4 induced by EphrinA1 exacerbates disruption of the blood-brain barrier following cerebral ischemia-reperfusion via the Rho/ROCK signaling pathway. Experimental and Therapeutic Medicine [Internet]. 2018;16:1–8. Available from: https://www.ncbi.nlm.nih.gov/pmc/articles/PMC6122430/pdf/etm-16-03-2651.pdf

68. Figueroa JD, Benton RL, Velazquez I, Torrado AI, Ortiz CM, Hernandez CM, et al. Inhibition of EphA7 up-regulation after spinal cord injury reduces apoptosis and promotes locomotor recovery. J Neurosci Res. 2006;84:1438–51.

69. Goldshmit Y, Spanevello MD, Tajouri S, Li L, Rogers F, Pearse M, et al. EphA4 Blockers Promote Axonal Regeneration and Functional Recovery Following Spinal Cord Injury in Mice. Plos One. 2011;6:e24636.

70. Bundesen LQ, Scheel TA, Bregman BS, Kromer LF. Ephrin-B2 and EphB2 Regulation of Astrocyte-Meningeal Fibroblast Interactions in Response to Spinal Cord Lesions in Adult Rats. J Neurosci. 2003;23:7789–800.

71. Jacobi A, Schmalz A, Bareyre FM. Abundant Expression of Guidance and Synaptogenic Molecules in the Injured Spinal Cord. Plos One. 2014;9:e88449.

72. Dvoriantchikova G, Pappas S, Luo X, Ribeiro M, Danek D, Pelaez D, et al. Virally delivered, constitutively active NFκB improves survival of injured retinal ganglion cells. Eur J Neurosci. 2016;44:2935–43.

73. Tameling C, Stoldt S, Stephan T, Naas J, Jakobs S, Munk A. Colocalization for super-resolution microscopy via optimal transport. Nat Comput Sci. 2021;1:199–211.

74. Strat AN, Kirschner A, Yoo H, Singh A, Bagué T, Li H, et al. Engineering a 3D hydrogel system to study optic nerve head astrocyte morphology and behavior. Exp Eye Res. 2022;220:109102.

75. Saha S, Greferath U, Vessey KA, Grayden DB, Burkitt AN, Fletcher EL. Changes in ganglion cells during retinal degeneration. Neuroscience. 2016;329:1–11.

76. Wan Y, Yang J-S, Xu L-C, Huang X-J, Wang W, Xie M-J. Roles of Eph/ephrin bidirectional signaling during injury and recovery of the central nervous system. Neural Regen Res. 2018;13:1313–21.

77. Li L, Huang H, Fang F, Liu L, Sun Y, Hu Y. Longitudinal Morphological and Functional Assessment of RGC Neurodegeneration After Optic Nerve Crush in Mouse. Front Cell Neurosci. 2020;14:109.

78. Feng G, Mellor RH, Bernstein M, Keller-Peck C, Nguyen QT, Wallace M, et al. Imaging Neuronal Subsets in Transgenic Mice Expressing Multiple Spectral Variants of GFP. Neuron. 2000;28:41–51.

79. Blandford SN, Hooper ML, Yabana T, Chauhan BC, Baldridge WH, Farrell SRM. Retinal Characterization of the Thy1-GCaMP3 Transgenic Mouse Line After Optic Nerve Transection. Investigative Opthalmology Vis Sci. 2019;60:183.

80. Hassan-Mohamed I, Giorgio C, Incerti M, Russo S, Pala D, Pasquale EB, et al. UniPR129 is a competitive small molecule Eph-ephrin antagonist blocking in vitro angiogenesis at low micromolar concentrations. Brit J Pharmacol. 2014;171:5195–208.

81. Martiny-Baron G, Holzer P, Billy E, Schnell C, Brueggen J, Ferretti M, et al. The small molecule specific EphB4 kinase inhibitor NVP-BHG712 inhibits VEGF driven angiogenesis. Angiogenesis. 2010;13:259–67.

82. Jellinger KA. Basic mechanisms of neurodegeneration: a critical update. J Cell Mol Med. 2010;14:457–87.

83. Sheikh S, Safia, Haque E, Mir SS. Neurodegenerative Diseases: Multifactorial Conformational Diseases and Their Therapeutic Interventions. J Neurodegener Dis. 2013;2013:563481.

84. Wareham LK, Liddelow SA, Temple S, Benowitz LI, Polo AD, Wellington C, et al. Solving neurodegeneration: common mechanisms and strategies for new treatments. Mol Neurodegener. 2022;17:23.

85. Tuttle R, Braisted JE, Richards LJ, O’Leary DD. Retinal axon guidance by region-specific cues in diencephalon. Dev Camb Engl. 1998;125:791–801.

86. Tadesse T, Cheng Q, Xu M, Baro DJ, Young LJ, Pallas SL. Regulation of ephrin-A expression in compressed retinocollicular maps. Dev Neurobiol. 2013;73:274–96.

87. Cheng H-J, Nakamoto M, Bergemann AD, Flanagan JG. Complementary gradients in expression and binding of ELF-1 and Mek4 in development of the topographic retinotectal projection map. Cell. 1995;82:371–81.

88. Yang J-S, Wei H-X, Chen P-P, Wu G. Roles of Eph/ephrin bidirectional signaling in central nervous system injury and recovery. Experimental and Therapeutic Medicine [Internet]. 2018;15:1–9. Available from: https://www.ncbi.nlm.nih.gov/pmc/articles/PMC5795627/pdf/etm-15-03-2219.pdf

89. Joly S, Jordi N, Schwab ME, Pernet V. The Ephrin receptor EphA4 restricts axonal sprouting and enhances branching in the injured mouse optic nerve. Eur J Neurosci. 2014;40:3021–31.

90. Vilallongue N, Schaeffer J, Hesse A-M, Delpech C, Blot B, Paccard A, et al. Guidance landscapes unveiled by quantitative proteomics to control reinnervation in adult visual system. Nat Commun. 2022;13:6040.

91. Benedetto BD. Ephrins in astrocytes: synaptic erasers on stage. Oncotarget. 2017;5:5676–7.

92. Heneka MT, Carson MJ, Khoury JE, Landreth GE, Brosseron F, Feinstein DL, et al. Neuroinflammation in Alzheimer’s disease. Lancet Neurology. 2015;14:388–405.

93. Lopez-Rodriguez AB, Hennessy E, Murray CL, Nazmi A, Delaney HJ, Healy D, et al. Acute systemic inflammation exacerbates neuroinflammation in Alzheimer’s disease: IL- 1β drives amplified responses in primed astrocytes and neuronal network dysfunction. Alzheimer’s Dementia. 2021;17:1735–55.

94. Wang H, Edwards G, Garzon C, Piqueras C, Bhattacharya SK. Aqueous humor phospholipids of DBA/2J and DBA/2J-Gpnmb +/SjJ mice. Biochimie. 2015;113:59–68.

95. Himanen JP, Yermekbayeva L, Janes PW, Walker JR, Xu K, Atapattu L, et al. Architecture of Eph receptor clusters. Proceedings of the National Academy of Sciences of the United States of America. 2010;107:10860–5.

96. Kania A, Klein R. Mechanisms of ephrin–Eph signalling in development, physiology and disease. Nat Rev Mol Cell Bio. 2016;17:240–56.

97. Miller NR, Arnold AC. Current concepts in the diagnosis, pathogenesis and management of nonarteritic anterior ischaemic optic neuropathy. Eye. 2015;29:65–79.

98. Alves JM, Seabra M, Braz L, Guimarães J. Optic neuropathy: A 15-year retrospective observational study. Mult Scler Relat Dis. 2020;44:102337.

99. Siyanaki MRH, Azab MA, Lucke-Wold B. Traumatic Optic Neuropathy: Update on Management. Encycl. 2023;3:88–101.

100. Tezel G, Thornton IL, Tong MG, Luo C, Yang X, Cai J, et al. Immunoproteomic Analysis of Potential Serum Biomarker Candidates in Human Glaucoma. Invest Ophth Vis Sci. 2012;53:8222–31.

